# High throughput steady-state enzyme kinetics measured in a parallel droplet generation and absorbance detection platform

**DOI:** 10.1101/2022.07.28.500969

**Authors:** Stefanie Neun, Liisa van Vliet, Florian Hollfelder, Fabrice Gielen

## Abstract

Microfluidic water-in-oil emulsion droplets are becoming a mainstay of experimental biology, where they replace the classical test tube. In most applications (e.g. in ultrahigh throughput directed evolution) the droplet content is identical for all compartmentalized assay reactions. When emulsion droplets are used for kinetics or other functional assays, though, concentration dependencies (e.g. of initial rates for Michaelis-Menten plots) are required. Droplet-on-demand systems satisfy this need but extracting large amounts of data is challenging. Here we introduce a multiplexed droplet absorbance detector which, coupled to semi-automated droplet generation, forms a tubing-based droplet-on-demand system able to generate and extract quantitative datasets from defined concentration gradients across multiple series of droplets for multiple time points. The emergence of product is detected by reading the absorbance of the droplet sets at multiple, adjustable time points (reversing the flow direction after each detection, so that the droplets pass a line scan camera multiple times). Detection multiplexing allows absorbance values at twelve distinct positions to be measured and enzyme kinetics are recorded for label-free concentration gradients (composed of about 60 droplets each, covering as many concentrations). With a throughput of around 8640 data points per hour, a 10-fold improvement compared to the previously reported single point detection method is achieved. In a single experiment, twelve full datasets of high-resolution and high accuracy Michaelis-Menten kinetics were determined to demonstrate the potential for enzyme characterization for glycosidase substrates covering a range in enzymatic hydrolysis of seven orders of magnitude in k_cat_/K_M_. The straightforward set-up, high throughput, excellent data quality, wide dynamic range that allows coverage of diverse activities suggest that this system may serve as a miniaturized spectrophotometer to for detailed analysis of study clones emerging from large-scale combinatorial experiments.

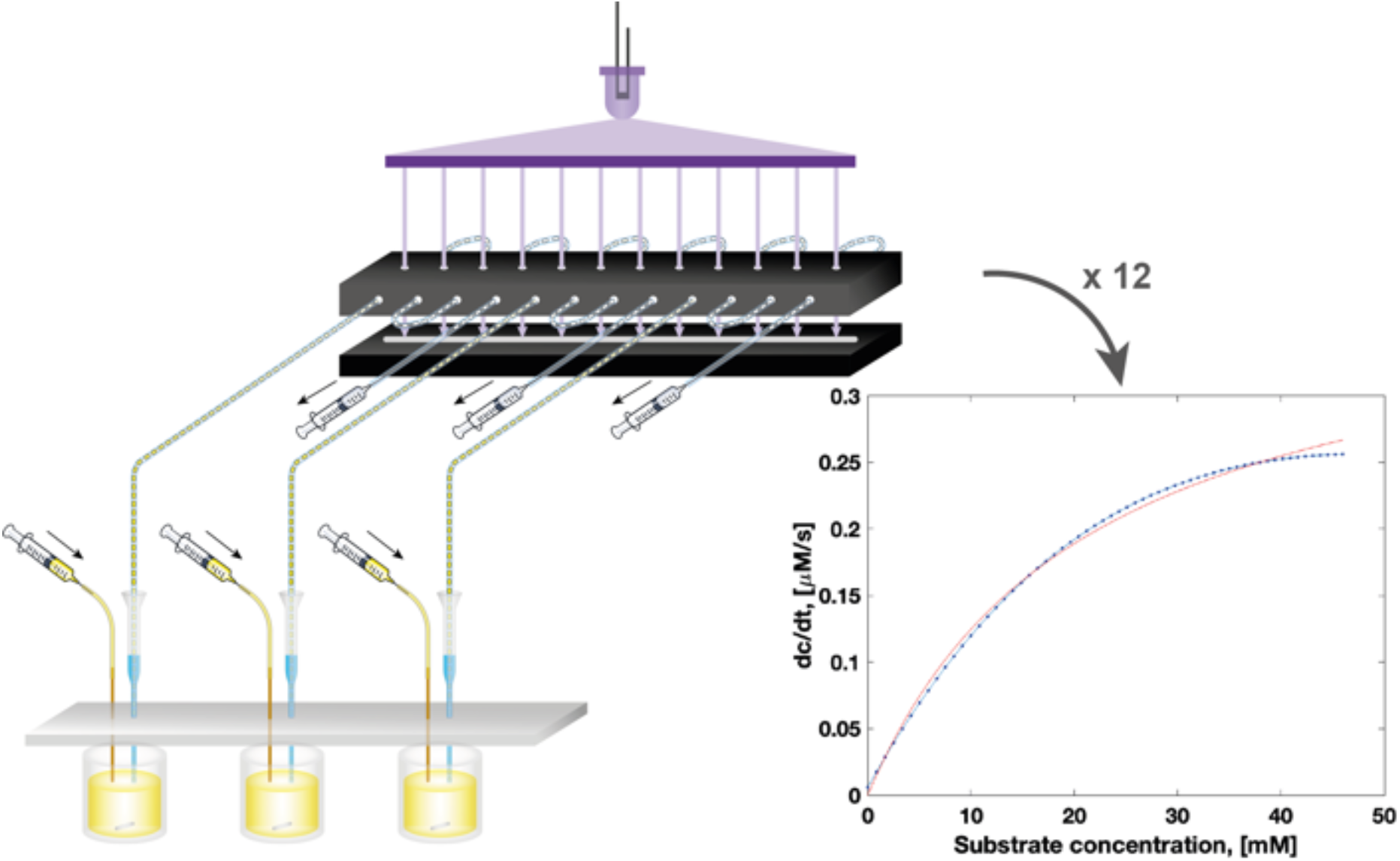

## INTRODUCTION

Enzymes hold tremendous potential as efficient and sustainable biocatalysts for applications ranging from energy generation to recycling. Concomitant with the decline in the cost of next-generation sequencing and the emergence of large-scale metagenomic sequencing projects^1-2^ we have gained access to protein sequence information at rates never before seen. The MGnify database alone currently holds more than two billion non-redundant protein sequences from microbiomes and keeps growing exponentially.^3^ To unearth the treasures contained in this trove, however, experimental testing of candidate enzymes is necessary, even if the bioinformatic analysis or structure prediction by AlphaFold2^4^ can reduce their number, because unambiguous extrapolation to function from these lines of evidence is not yet possible. In fact, the number of experimentally characterized proteins is small compared to the available sequence information,^5^ so prediction of new, hypothetical gene functions is hard, making further experimental characterization necessary.

Extensive experimental characterization is also a prerequisite for a mechanistic analysis, especially when extended interaction networks in active sites, described as sectors or units,^6-8^ operate with functional synergy of amino acids. Likewise, the analysis of intra-gene epistasis in protein evolution trajectories^9-11^ relies on cooperative effects of amino acids. To resolve such questions, the analysis of single mutants, e.g. alanine-knock-outs of active site residues, is insufficient. The numbers arising from combinations of mutants are, of course, much larger and their quantitative characterization requires a scale-up of assays systems. For these enzymological challenges multi-well plate-based measurements can become uneconomical and unwieldy, pointing towards miniaturized systems as alternatives. Progress in the development of microfluidic tools^12-18^ for the functional screening of enzyme libraries at ultrahigh throughput now allows us to routinely screen millions of enzyme variants and identify active variants in directed evolution and functional metagenomic projects.^19-22^ By contrast, the methods used to profile newly discovered enzymes with kinetic studies remain mostly unchanged, requiring a large number of tests for full and quantitative enzyme characterization, including steady-state or pre-steady-state kinetic assays, thermostability assessments, substrate specificity and enzymatic activity under different experimental conditions (e.g. in the presence of additives and/or inhibitors).

For the kinetic characterization of enzyme activity initial reaction velocities need to be recorded across a large range of substrate concentrations to define a non-linear Michaelis-Menten plot. When carried out in a standard lab, this process entails tedious pipetting work and can be costly, if the reagents involved are expensive or hard to synthesize. The pipetting necessary to set up concentration gradient can be substituted by liquid-handling robots, but only a 2-3-fold acceleration in comparison to manual handling is expected^23^ and a lot of plasticware is necessary. Moreover, traditional techniques typically rely on 96- or 384-well plates with working volumes of 200 μL or 50 μL per reaction condition, respectively and therefore, require large amounts of enzyme and substrate. Thus a number of approaches have been devised to find practical solutions for miniaturization of kinetic measurements in order to increase the assay throughput and reduce the required reagent volumes.^8,24-30^ However, many of these systems (Tables S1-3, Supporting Information) are costly and difficult to implement. Real world uptake (e.g. in a standard molecular biology lab) leading to real improvements in throughput is still an unresolved challenge.

We have previously introduced an inexpensive droplet-on-demand microfluidic system for the generation of droplets with defined contents, e.g. setting up accurate concentration gradients for the determination of enzyme kinetics based on an absorbance readout.^31^ Here, we developed a multiplexed absorbance reader for measuring reaction progress in series of droplets stored in tubing to increase the throughput of our droplet-on-demand platform.^31^ The parallelized monitoring of multiple enzymatic reactions in nanoliter droplets is implemented in a line camera detection scheme, with which the absorbance values at twelve distinct positions can be measured. Three series of droplets with label-free concentration gradients (composed of about 60 droplets each) are generated in parallel and their absorbance is recorded with a throughput of around 8640 data points per hour (Figure 1), representing a 10-fold improvement in throughput.^28^ In a single experiment, twelve full datasets of high-resolution and high accuracy Michaelis-Menten kinetics were determined, in parallel for three different substrates for a promiscuous metagenome-derived glycosidase. To demonstrate the potential of our platform as a generic enzyme characterization tool, we also detail the detection limits to quantify both efficient and inefficient enzyme catalysis with a range of glycosidase substrates, highlighting the relevance for the assessment of promiscuous activities.

**Figure 1.**
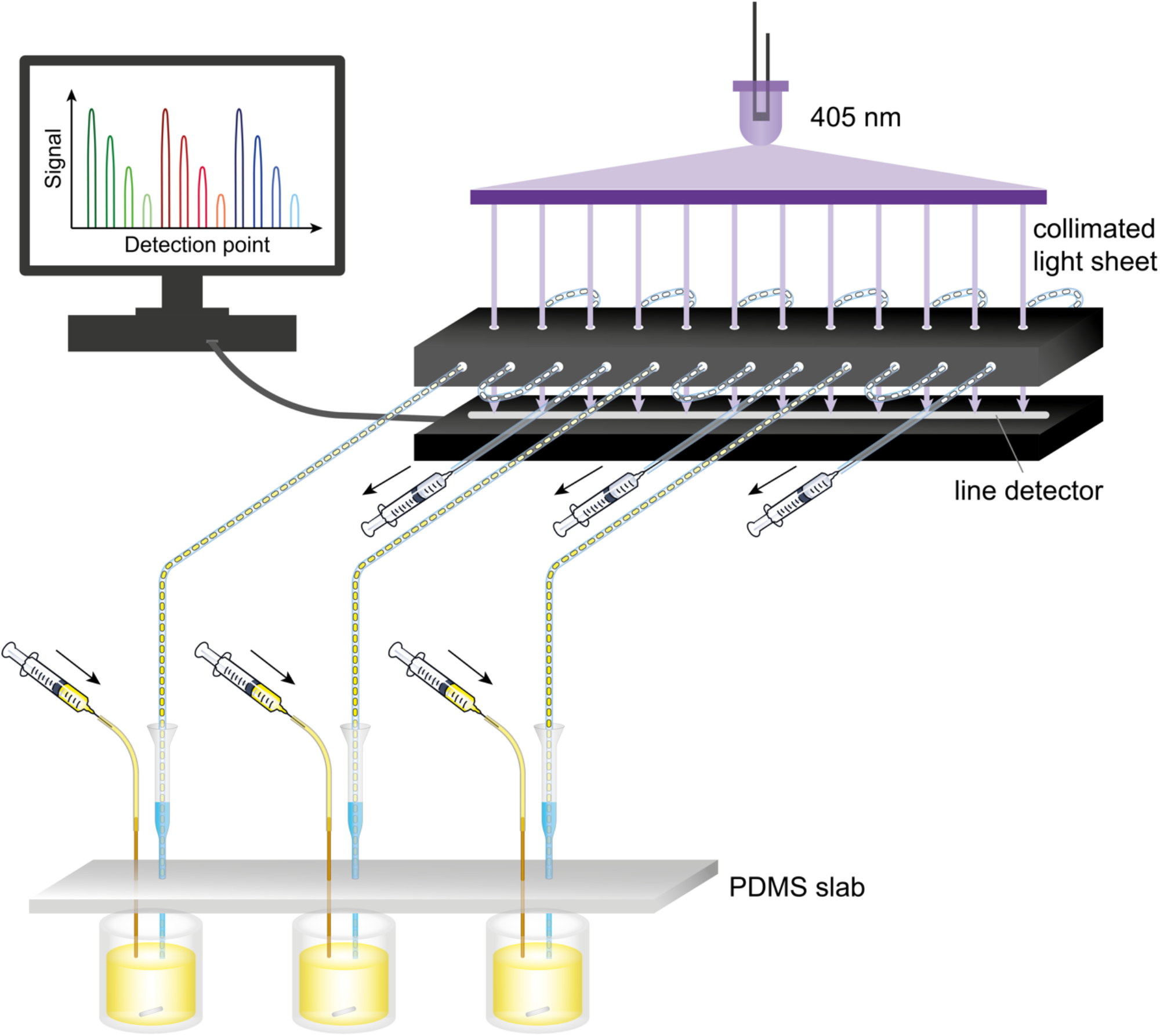
Schematic overview of the microfluidic platform. Droplets are made from three enzyme solutions in reaction vessels in parallel (see Figure 2a) by pulling liquid with a syringe from the end of each tubing. Tubing is held in place vertically by a PDMS slab (pictured in gray) sitting on top of a 384-well plate. Through injection of a substrate solution into the reaction well and fast mixing with a magnetic stirrer, a gradient of defined concentrations in droplets is generated. Each line passes four times through the collimated light sheet (optical setup for its generation in Figure 2b) and the absorbance at 405 nm of the droplets is recorded by a line camera below to generate four separate datasets per droplet gradient. To read the absorbance of the droplet sets at multiple time points the flow direction is switched between pulling and pushing mode.

## METHODS

### Optics

The light steering module producing a collimated light line was assembled from parts sourced from Thorlabs: a fiber-coupled LED (405 nm, M405FP1), an LED controller (LEDD1B), an SMA-to-SMA multimode fiber optic with core diameter 1 mm (M53L), a fiber collimator (CVH100-COL), a line diffuser (ED1-L4100-MD), and a light collimator (SM2F). The line beam was projected onto a CCD Line Camera (LC100, 2048 pixels, detector range: 350-1100 nm, pixel size 14 × 56 µm, 14 µm pitch).

### Fluidic connector

The fluidic connector aligning tubing with the light sheet at 12 separate positions was designed on a CAD software (DraftSight) and fabricated by laser cutting (Razorlabs) in black acrylic. The individual parts were assembled with epoxy glue.

### Droplet gradient generator

Each droplet generator was custom-made from the narrow part cut from a 200 µl round gel-loading tip (Starlab) and a yellow pipette tip for 2-200 μL pipettes (200 μL TipOne tip, Starlab). These pipette tip parts were connected through a hole in a PDMS slab with the narrower tip below and the wider yellow tip above the PDMS. PTFE tubing (ultramicrobore, 0.2 mm ID, 0.36 mm OD; Cole-Parmer) was inserted into the tips (to end just about 1 mm inside from the end of the gel-loading tip). The space between the tip wall and the tubing was filled with 20 μL HFE-7500 containing 0.1% 008-Fluorosurfactant (RAN Biotechnologies) and 35% 1-bromo-3,5-bis(trifluoromethyl)benzene (Merck). Each PTFE tubing was threaded four times through the fluidic connector and fused to wider PE tubing (0.38 mm ID, 1.09 mm OD; Portex) connected to a 100 μL gas-tight glass syringe (SGE). The tubing and syringe (to about 60 μL syringe volume) were filled with the oil specified above. The substrate injector was made from fused silica tubing (200 μm ID, 360 μm OD; Polymicro Technologies) conjoint with PE tubing (0.38 mm ID, 1.09 mm OD; Portex), which was connected to a 100 μL glass syringe (SGE). The glass syringe and tubing were filled with deionized water followed by a small plug of air before the substrate solution (exactly 7 μL) was loaded into the tubing. This substrate injector was inserted into the reaction well through a separate hole in the PDMS slab.

### Formation of droplet gradients

A 384-well plate was placed on a magnetic stirrer (IKA). For each droplet generator, one well of the plate was filled with 40 μL enzyme solution and a magnetic stir bar was placed inside. The neighboring well contained 60 μL of the oil mixture and a third well (one row below) was filled with 60 μL 1 mM *p*NP solution in the reaction buffer. Before starting droplet generation from the reaction solution, a few (∼5) air bubbles were made to confirm proper device operation. Subsequently, droplets were made by pulling liquid into the droplet generator at 4 μL/min, while injecting 5 μL substrate solution at a rate of 10 μL/min under magnetic stirring at 1500 rpm. This generated a concentration gradient distributed over approximately 60 droplets for 30 s. After droplet generation was complete, the flow was shortly paused, and the droplet generators were manually moved to the oil containing wells to continue pulling oil without making additional droplets. Once droplets had passed through all measurement points, the flow direction was inverted to push the droplets at 4 μL/min. The flow directions were regularly alternated over 30 min. Before switching to the last round in pushing mode, droplets were generated from the 1 mM *p*NP solution and measured to calibrate the absorbance values for each detection point.

### Enzymatic reactions

SN243 was expressed and purified as described in Neun et al. ^22^ Kinetics were determined in 100 mM Tris pH 8.0, 150 mM NaCl at room temperature. For kinetic measurements in droplets, the following concentrations were used: injection of 500 μM *p*NP-β-D-glucuronide (*p*NP-β-GlcA) into 5 nM SN243; 100 mM *p*NP-β-D-galacturonide (*p*NP-β-GalA) into 25 nM SN243; 400 mM *p*NP-β-D-glucopyranoside (*p*NP-β-Glc) or *p*NP-β-D-xylopyranoside (*p*NP-β-Xyl) into 1 μM SN243; 400 mM *p*NP-β-D-galactopyranoside (*p*NP-β-Gal) or *p*NP-α-L-arabinofuranoside (*p*NP-α-Ara*f*) into 10 μM SN243. All substrates were dissolved in DMSO.

### Software

A custom-made Labview script (https://github.com/fhlab/Line_detector_kinetics) automated acquisition at 200 Hz of transmitted light (area under the curve) for 12 positions manually set and stored the time and raw signal data in a single .csv file.

## RESULTS AND DISCUSSION

### Design of a parallel droplet generation and absorbance detection platform

We designed and built a parallel droplet sampling microtiter plate adaptor with an absorbance detector unit to accelerate enzyme characterization. Each droplet generation unit in our platform operates by negative pressure, as previously described in Gielen et al. (2015).^31^ Briefly, a pulseless syringe pump pulls liquid with high accuracy during droplet formation. Microbore polytetrafluoroethylene (PTFE) tubing is manually inserted into a tapered pipette tip whose ID matches the OD of the tubing. The pipette tip is cut closely to the end of the tubing and then the pipette tip is filled with carrier oil around the tubing. This system is placed into the reaction vessel (here a well of a 384-well plate) and, by aspiration, aqueous droplets break off separated by the carrier oil in regular intervals, ensuring the generation of monodisperse plugs with constant oil separation. Three such droplet makers were operated in parallel thanks to the design of a bespoke PDMS slab adaptor keeping droplet making and substrate injection tubing aligned with each well (Figure 1 and Figure 2a). Droplets were formed by a multi-rack pump connected to three syringes and operating in withdrawal mode producing the droplets while another syringe pump injected three substrate solutions through three separate tubing connections. In this way, droplet makers could be conveniently moved to other wells, allowing us to load gradients sequentially, space them out with more oil or add calibration droplets. Next, we have designed and built a parallelized droplet absorbance detector enabling measurements at up to 12 separate locations. This detection scheme not only increases data acquisition rates but provides an opportunity to measure a given reaction multiple times at arbitrary time intervals.

**Figure 2.**
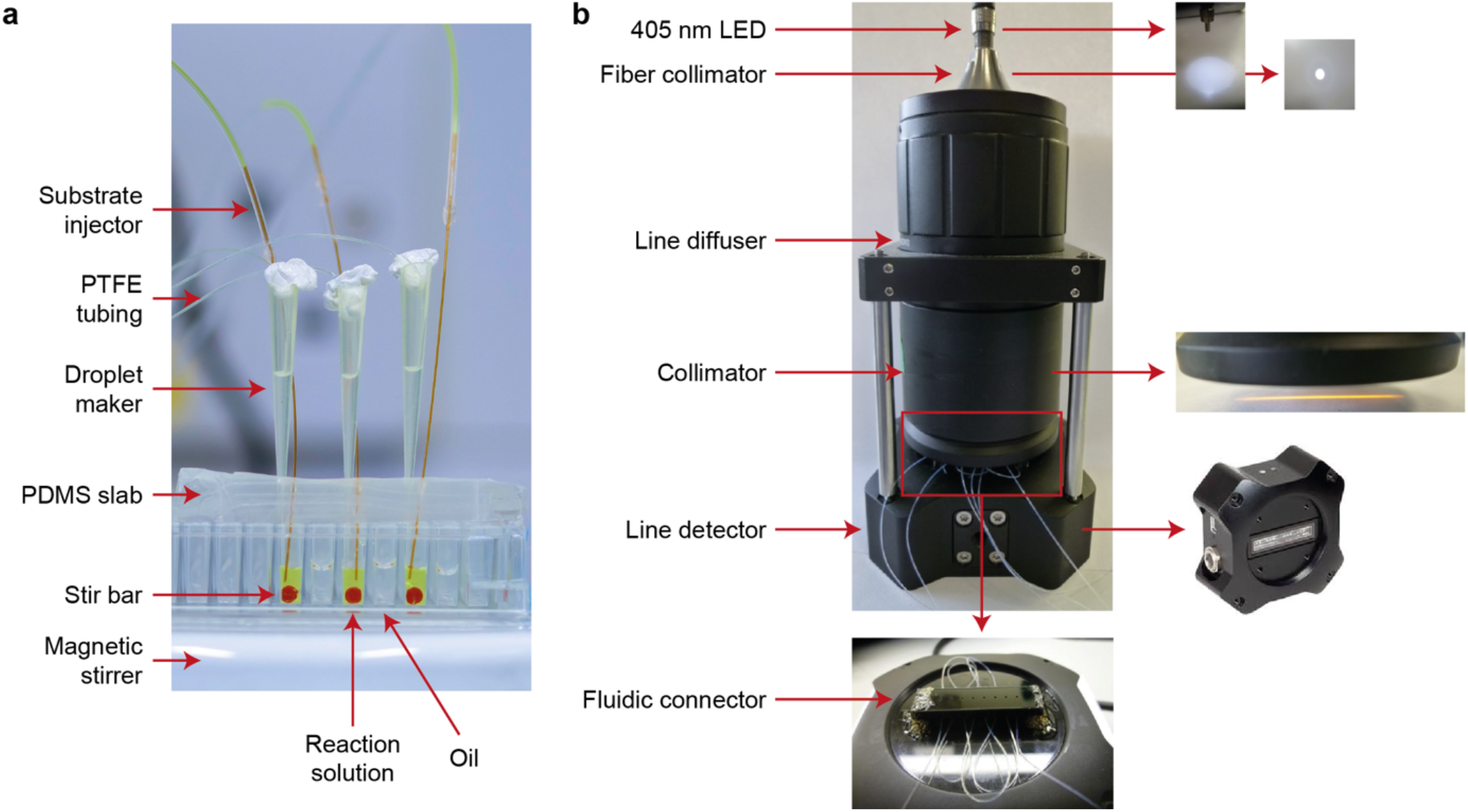
Setup of the microfluidic platform for the generation of droplet gradients and optical detection by a line camera. **(a)** Photograph of a set of three droplet generators operated in parallel. After droplet formation, the droplet generators are all moved one well further to the right containing oil to stop making droplets and pull oil instead. **(b)** Schematic of the parallelized detection setup. The optical setup generates a collimated light sheet which illuminates all detection points of a line camera uniformly. Droplets produced from individual droplet generators **(a)** converge towards the line camera where their absorbance is read. A more detailed view of the fluidic connector is shown in Figure S3 in the Supporting Information.

Briefly, the light steering elements produced a collimated line beam towards a CCD line scan camera, measuring changes in transmitted light intensity during droplet transit. Tubing containing droplets was placed in the light pass so that absorbance could be measured at specific locations (Figure 2b). The light steering module was pre-assembled and subsequently mounted onto the line camera using mounting rods. A fiber-coupled LED (405 nm) sent light through to a fiber collimator and then to a line diffuser that generated a homogeneous line pattern with low divergence. A second collimator directed the line illumination pattern towards a custom-made connector whose function was to align the tubing through which droplets were shuttled with the line pattern. The connector was made of black acrylic to prevent transmissions and reflections. It possessed 12 vertical holes of 400 ± 10 µm diameter in two aligned parallel plates located above and below the height at which tubing was placed. The tubing containing the droplets was aligned with the 12 horizontal holes using 2 vertical plates also having 12 aligned holes (Figure S3, Supporting Information). This ensured that most of the light rays traveled only through the microbore tubing, reducing background signal and blocking any scattered light. The tubing was manually threaded through the connector before being placed onto the line camera. The camera collected and recorded the light intensities at the 12 separate locations equally spaced over a length of 5 cm. This configuration allowed for rapid interchange of tubing when starting new experiments and fine adjustment of tubing insertion points to control the time interval in between measurements.

Initially, we verified that no stray light coming from the gap between the line camera and the collimator was detected. Next, the LED light intensity was manually adjusted so that, in absence of any droplets, the detection point receiving the highest intensity would still be below the saturation of the detector. The uniformity of illumination across the points was checked first in the presence of inserted tubing and showed good homogeneity across the line camera: we observed a maximum difference of ∼30% deviation from the maximum light transmission. We assign this difference to the tolerance in the fabrication of the holes leading to imprecise alignment of tubing. However, this range was found to be reproducible across separate experiments. After insertion of tubing, we calibrated every detection point by measuring a model gradient of *para*-nitrophenol (*p*NP). This calibration translated into differences for limits of detection (LODs) ranging from 5 μM to 18 µM *p*NP with a typical mean of 10 μM with a 95% confidence level (Figure S2, Supporting Information). Although the LOD was found to be higher than in our previously reported technique (3 μM) using a photodetector with a wide photon collection area, it was not far off for the most sensitive points.^28^

In comparison to implementations using individual LEDs and detectors for every position,^27, 32^ our line scan detection system avoids separate calibration of multiple LEDs. The system is much easier to assemble, using a single light source, and generates data from a single line camera while retaining high UV-Vis sensitivity. Moreover, to accommodate for the differences in absorbance spectra of chromophores, the LED can be easily exchanged to one with the best suited wavelength as the thin walls of the PTFE capillary have a low baseline absorption in the UV-Vis spectrum.

Here, we use a 405 nM LED to detect *p*NP, a commonly used leaving group in commercial model substrates for hydrolase reactions. This leaving group is not suited for classic microfluidic screening formats, because of its tendency to leak into the oil phase, but the confined droplet format enables the reliable detection of *p*NP in droplets: the use of considerably large droplets (∼30 nL), which implies a small surface to volume ratio, and a very low surfactant concentration (0.1% 008-FluoroSurfactant, RAN Biotechnologies) slow the leakage process down substantially as droplets rarely get in contact with each other. Moreover, due to the fast generation of droplets, detection on the same platform and short assay times needed (<30 minutes), inter-droplet transport of *p*NP was not observed in our setup.

In our microfluidic platform three droplet generators operate in parallel, feeding four detection points each by running the tubing in a serpentine loop through the holes of a bespoke fluidic connector placed over the line camera (Figure 1). It would be possible to increase the number of droplet generation systems in the platform further to up to twelve, so that each of them feeds into one detection point (see Figure S4, Supporting Information). The absorbance of droplets is monitored by a Labview script, which calculates (at a sampling rate of 200 Hz) the area under the curve as a measure for the total light intensity between two fixed positions corresponding to each of the 12 holes. To avoid large fluctuations in measured absorbance values observed at the droplet edges (that result from a refractive index mismatch between the aqueous and fluorous oil phase and complicate data analysis), the carrier oil (HFE-7500 supplemented with 0.1% 008-FluoroSurfactant) contained additionally 35% 1-bromo-3,5-bis(trifluoromethyl)benzene for refractive index matching.^33^

### Generation of concentration gradients in droplets and detection across all twelve points

In order to determine enzyme kinetics according to Michaelis-Menten, the initial velocities (v_0_) of the catalyzed reaction need to be measured across a wide range of substrate concentrations. In analogy to a microtiter plate setup, where every condition needs to be optically followed in a separate well, droplets serve as reaction compartments and each droplet contains a different amount of substrate. We generate the gradients by injecting a defined volume of the substrate (5 μL) at a specified flow rate (10 μL/min) into the microtiter plate well containing the enzyme solution (40 μL) while fast mixing with a magnetic stirrer and generating droplets. To first check how the product concentration in droplets translates into absorbance (A_405_) in our system, we injected 20 mM *p*NP into reaction buffer and recorded A_405_ across all twelve points. The calibration curves obtained indicated linear correlation between detector response and concentration of *p*NP up to above 1 mM *p*NP (Section S2, Supporting Information). The initial slopes measured in our setup never reached such high product levels so that quantification of absolute *p*NP per droplet could be accurately calculated using a linear correlation.

### Highly accurate determination of enzyme kinetics in microfluidic droplets

To validate our system, we compared standard well plate datasets to kinetic parameters obtained in the droplet format. A substrate gradient across ∼60 droplets was generated and v_0_ determined for 60 distinct substrate concentrations simultaneously. In order to obtain v_0_, measurements at multiple time points are required. Once droplets have passed through their respective detection points, inverting the flow direction of the system by switching between pulling and pushing modes of the syringe pump allowed us to optically follow the reaction over time. In the last pulling phase, droplets with 1 mM *p*NP were generated and their absorbance was detected in order to later calculate the product concentrations in droplets from the recorded optical signal. Each droplet is defined by its position in the gradient and receives an accurate timestamp for every detected signal. Therefore, extraction of (a) the substrate concentration of each droplet by its position within the gradient, (b) the absorbance information for a specific droplet at every time it passed through the detection point, and (c) the calibration of a specific point for its linear dependence of optical signal to product concentration allowed for the determination of v_0_ for each substrate concentration. Combining this information for all droplets detected in one point, yields a dataset with which Michaelis-Menten kinetics could be approximated with very high accuracy.

In Figure 3, we show and explain the step-by-step generation of kinetic data from one detection point for the reaction of SN243 with *p*NP-β-Xyl (see also Section S1, Supporting Information). The values determined for k_cat_ (0.4 s^-1^) and K_M_ (21.3 mM) differ only minimally from the kinetic data obtained in microtiter plate (k_cat_ = 0.4 s^-1^, K_M_ = 23.0 mM).^22^ Comparison of the datasets from all twelve detection points (Figures S5-S12, Supporting Information), which were obtained from three different sets of droplets (each feeding four detection points) proved reliable reproducibility of the data. Mean values and their standard deviation averaged from the twelve datasets for the hydrolysis of *p*NP-β-Xyl resulted in k_cat_ = 0.4 ± 0.04 s^-1^ and K_M_ = 21.4 ± 2.2 mM with the highest relative difference of 10% in k_cat_ and 15% in K_M_. Considering the high number of individual concentrations for which v_0_ was determined, the production of three separate sets of droplets, and the measurement of four full kinetic datasets for each set of droplets, it could be argued that the Michaelis-Menten parameters obtained from microfluidic droplets is better supported, based on a 50-fold larger dataset compared to a typical plate reader experiment and is providing more accurate approximations.

**Figure 3.**
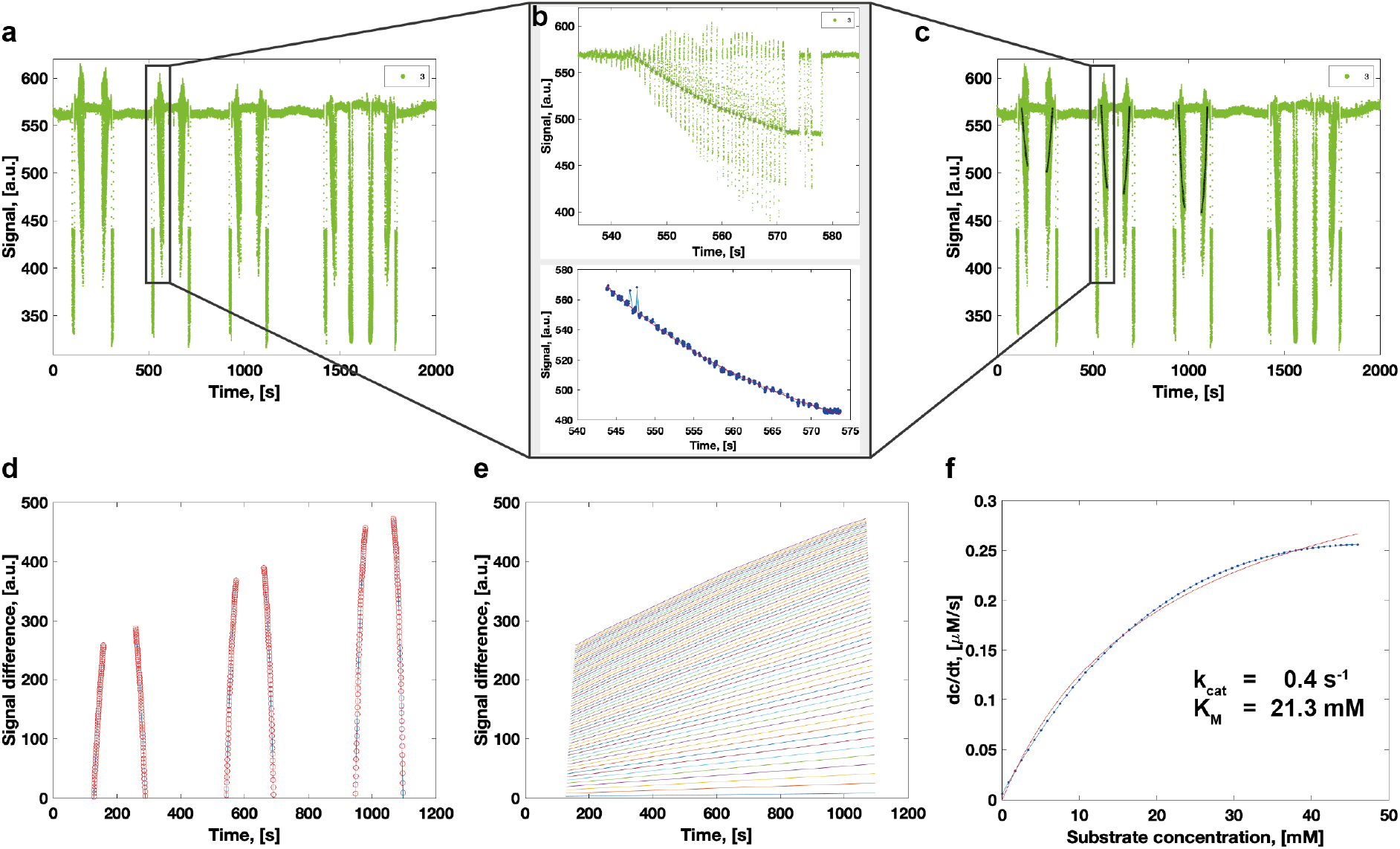
Example analysis for the approximation of Michaelis-Menten kinetics from one detection point of the microfluidic platform. The example shows data recorded for the hydrolytic cleavage of *p*NP-β-Xyl by SN243. **(a)** Raw signal recorded for the absorbance at 405 nm. The first six measurements of the droplet gradient were used to determine the kinetic parameters. **(b)** Zoom into the third absorbance recording for the droplet gradient (top) and filtering to the signal corresponding to aqueous phase in droplets (bottom). **(c)** Recorded signal overlaid with the identified and fitted concentration gradients. **(d)** Signal difference to the enzyme-only baseline for the individual droplets with increasing substrate concentration of the gradient at the six time points. **(e)** Signal difference for each droplet (i.e. substrate concentration) vs. time for the determination of initial reaction velocities. **(f)** Approximation of the initial reaction velocities (dc/dt) to the Michaelis-Menten equation.

### Wide dynamic range for the determination of kinetic parameters

To investigate whether our platform also performs well for the determination of kinetic parameters of enzymatic reactions with a range of catalytic efficiencies and to test the detection limits of our system, we chose to measure Michaelis-Menten kinetics for the remaining five *p*NP-coupled glycoside substrates on which SN243 has recently been shown to be active.^22^ The kinetic parameters range from very efficient catalysis (k_cat_ = 23 s^-1^, K_M_ = 14 μM, with *p*NP-β-GlcA) to extremely low promiscuous activities (k_cat_ = 0.01 s^-1^, K_M_ = 85 mM, with *p*NP-α-Ara*f*), with a 10^7^-fold lower k_cat_/K_M_. We determined the kinetic parameters for SN243 with *p*NP-β-GlcA, *p*NP-β-GalA, *p*NP-β-Glc, *p*NP-β-Gal, and *p*NP-α-Ara*f* in the microfluidic platform and obtained similar values to those determined in the plate reader for all substrates (Table 1).

**Table 1:** Comparison of Michaelis-Menten parameters obtained from kinetic measurements in droplets and in microtiter plate for the hydrolytic cleavage of different *p*NP-glycoside substrates by SN243. Indicated values for parameters determined in droplet experiments are the mean values and their standard deviations averaged from twelve kinetic datasets in separate detection points. For Michaelis-Menten kinetics obtained from plate measurements, parameters are indicated for the fit of one dataset per reaction as well as standard errors with a 95% confidence interval. Kinetic data from plate reader measurements were taken from Neun et al.^22^

| Substrate | $K_M$, [mM] | | $k_{cat}$, [s<sup>-1</sup>] | | $k_{cat}/K_M$, [M<sup>-1</sup> s<sup>-1</sup>] | |
| --- | --- | --- | --- | --- | --- | --- |
|  | Droplets | Plate | Droplets | Plate | Droplets | Plate |
| pNP-β-GlcA* | $15.0 \times 10^{-3} \pm 3.6 \times 10^{-3}$ | $13.7 \times 10^{-3} \pm 0.9 \times 10^{-3}$ | $22.2 \pm 6.2$ | $23.1 \pm 0.2$ | $1.2 \times 10^6$ | $1.7 \times 10^6$ |
| pNP-β-Glc | $36.8 \pm 10.4$ | $56.5 \pm 4.1$ | $0.7 \pm 0.1$ | $0.7 \pm 0.03$ | $1.9 \times 10^1$ | $1.3 \times 10^1$ |
| pNP-β-GalA | $1.6 \pm 0.3$ | $1.0 \pm 0.1$ | $13.6 \pm 2.1$ | $16.6 \pm 0.7$ | $8.2 \times 10^3$ | $1.6 \times 10^4$ |
| pNP-β-Gal | $75.8 \pm 10.9$ | $38.7 \pm 4.6$ | $3.6 \times 10^{-2} \pm 0.5 \times 10^{-2}$ | $1.2 \times 10^{-2} \pm 0.1 \times 10^{-2}$ | $4.7 \times 10^{-1}$ | $3.1 \times 10^{-1}$ |
| pNP-β-Xyl | $21.9 \pm 1.6$ | $23.0 \pm 0.9$ | $0.4 \pm 0.02$ | $0.4 \pm 0.01$ | $1.8 \times 10^1$ | $1.8 \times 10^1$ |
| pNP-α-Araf | $107.8 \pm 37.4$ | $84.5 \pm 9.3$ | $2.7 \times 10^{-2} \pm 0.7 \times 10^{-2}$ | $1.4 \times 10^{-2} \pm 0.1 \times 10^{-2}$ | $2.6 \times 10^{-1}$ | $1.7 \times 10^{-1}$ |

\*Droplet measurements averaged from points 1 to 4 only.

Indeed, for *p*NP-β-GlcA averaged from four detection points, we determined a mean k_cat_ at 20 s^-1^ and K_M_ at 16 μM, demonstrating high accuracy for fast reactions with high affinity of the enzyme for the substrate. Going further to determine enzyme kinetics with an even higher catalytic efficiency than SN243 with *p*NP-β-GlcA (k_cat_/K_M_ = 1.2 × 10^6^ M^-1^s^-1^), we extrapolate that faster reactions (higher k_cat_) can be measured as the reaction can be slowed down by decreasing the enzyme concentration. However, the determination of higher substrate affinities depends on the sensitivity of the optical detection: similar to the limitations of a plate reader, we expect that the determination of lower K_M_ values (e.g. around 1 μM) for *p*NP as leaving group would become less accurate, as the detection of product formation for substrate concentrations below the K_M_ will be very close to the optical detection limit. Importantly, the sensitivity of the system for product detection depends also on the optical properties of the product itself, i.e. a lower concentration of a chemical moiety can be detected when its extinction coefficient is higher (obeying Beer-Lambert law).

On the other hand, when measuring the catalytic parameters for very inefficient enzymatic reactions, we observed that the lower limits for the accurate determination of k_cat_/K_M_ mainly depend on the intrinsic properties of the reaction components, namely substrate solubility and enzyme stability. If the substrate is not soluble at or above the concentration of the predicted K_M_ of the investigated reaction, only values covering the initial part of the Michaelis-Menten curve can be detected and the extrapolated kinetic parameters have a large error. This limitation is identical to a plate reader format for kinetic measurements. Additionally, the detection of decreased rates does not seem to have any limitations imposed by the microfluidic platform: if required, the absorbance of droplets can be determined over longer times, as droplets can be stored in-line and no product leakage was observed. If required, the enzyme concentration can be increased to generate more product within the same time and overcome the detection threshold. However, it is crucial for the determination of slow reactions that the enzyme is stable for the entire time of the kinetic measurement.

### Reliable generation of twelve kinetic datasets for three different reactions in half an hour

In a final set of kinetic measurements, we challenged the performance of the system to produce kinetic datasets for reactions with large differences in kinetic efficiency simultaneously and with high throughput. To this end, we produced three droplet gradients for the enzymatic reaction of SN243 with *p*NP-β-GalA, *p*NP-β-Xyl, and *p*NP-β-Gal in parallel, and determined kinetic datasets in four detection points for each of the reactions (Figure 4). All twelve Michaelis-Menten kinetics reproduced with high accuracy the previously determined kinetic parameters (Table 2). Indeed, despite using a fresh substrate solution and a new batch of purified enzyme, all determined parameters were within a small margin (i.e. in the same order of magnitude for k_cat_/K_M_) of the data, which had been determined across all twelve detection points of the system, and the plate reader results. The combination of multiple droplet gradient generating systems with the absorbance recording in a line camera increased the throughput of kinetic characterization by an order of magnitude.

**Table 2.** Kinetic parameters determined in parallel for SN243 with three different substrates each detected across four detection points of the microfluidic platform.

| Substrate | DP* | $K_M$, [mM] | $k_{cat}$, [s<sup>-1</sup>] | $k_{cat}/K_M$, [M<sup>-1</sup> s<sup>-1</sup>] |
| --- | --- | --- | --- | --- |
| pNP-β-GalA | 1 | 1.2 | 16.3 | $1.4 \times 10^4$ |
| | 2 | 1.0 | 15.0 | $1.4 \times 10^4$ |
| | 3 | $7.4 \times 10^{-1}$ | 12.3 | $1.7 \times 10^4$ |
| | 4 | $7.5 \times 10^{-1}$ | 12.5 | $1.7 \times 10^4$ |
| pNP-β-Gal | 5 | 79.0 | $4.3 \times 10^{-2}$ | $5.5 \times 10^{-1}$ |
| | 6 | 79.6 | $5.2 \times 10^{-2}$ | $6.5 \times 10^{-1}$ |
| | 7 | 70.5 | $4.3 \times 10^{-2}$ | $6.1 \times 10^{-1}$ |
| | 8 | 80.8 | $4.4 \times 10^{-2}$ | $5.5 \times 10^{-1}$ |
| pNP-β-Xyl | 9 | 23.5 | $4.8 \times 10^{-1}$ | $2.0 \times 10^1$ |
| | 10 | 22.1 | $5.7 \times 10^{-1}$ | $2.6 \times 10^1$ |
| | 11 | 22.0 | $5.5 \times 10^{-1}$ | $2.5 \times 10^1$ |
| | 12 | 20.2 | $5.4 \times 10^{-1}$ | $2.7 \times 10^1$ |

\*DP: detection point

**Figure 4.**
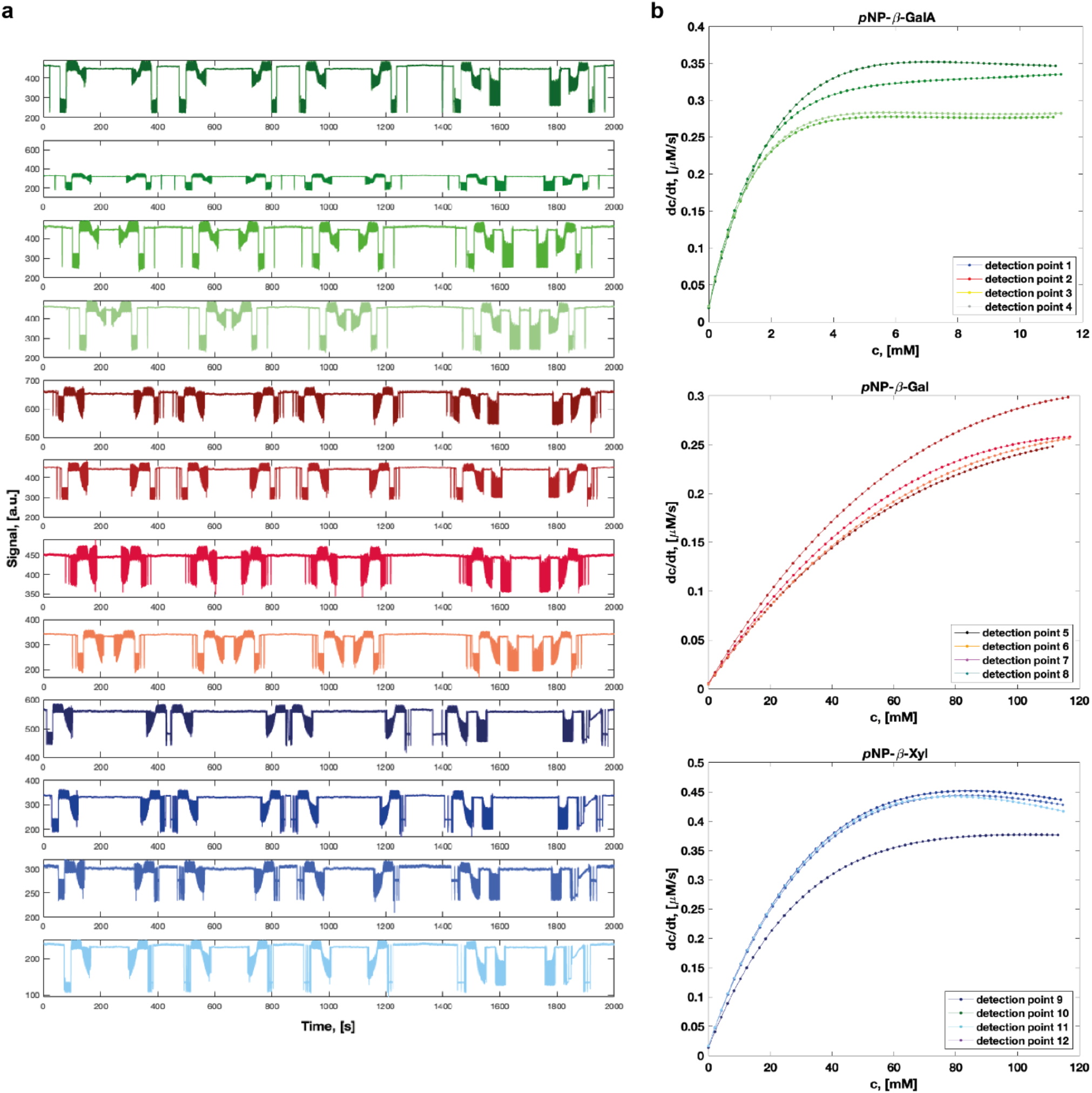
High-throughput determination of four full kinetic datasets for the hydrolysis of each of three different substrates by SN243 in parallel. **(a)** Raw signal for the twelve detection points with three different substrates. The reaction with *p*NP-β-GalA is detected successively in points 1-4, represented in the top four rows of the plot, *p*NP-β-Gal in 5-8, and *p*NP-β-Xyl in 9-12. **(b)** Initial velocities recorded at the twelve detection points are plotted against the substrate concentrations. Corresponding kinetic parameters are indicated in Table 2.

### High adaptability of the system to experimental requirements

While we demonstrated the use of the microfluidic platform to determine standard Michaelis-Menten kinetics, our platform offers flexibility to adapt to the specific requirements of a target reaction and to other applications. For instance, temperature control can be added to evaluate thermostability.^34^ In order to obtain more data points in shorter temporal increments, the length of the tubing between the droplet generation module and the first detection point, as well as in-between the detection points, can be reduced. If even higher temporal resolution is desired, it is possible to measure continuously the same droplet set across all twelve detection points within a single tubing and combine the data to a single kinetic dataset. This particular feature is enabled by the use of a CCD line camera and accurate calibration across all detection points. On the other hand, if a higher throughput with fewer data points is required, up to twelve reactions can be measured in parallel with one droplet generation module feeding one detection point. Moreover, similar to Michaelis-Menten kinetics measurements of initial rates can be used for kinetic characterization of inhibitors, as well as the determination of the optimal concentration of a reagent (e.g. co-factors) or buffer components (e.g. salt concentration in the buffer) for higher efficiency of the reaction. Further parallelization of the fluidic lines will enable probing of more than three enzymatic reactions at a time. For instance, we have built an adaptor interfacing eight separate wells to eight droplet making units (Figure S4, Supporting Information), highlighting the potential for direct interfacing with the classic microtiter plate format. Likewise, one could increase the number of measurement positions by reducing the distance between the holes of the fluidic connector. This will however be limited by crosstalk between the detection points that are too close to each other.

## CONCLUSIONS

Building on the ability to measure the dependence of enzymatic rate on substrate concentration in well-controlled gradients^31^ – with one droplet representing one concentration (‘droplet-on-demand’) – the platform presented in this study improves data interpretation and throughput. By generating continuous substrate gradients full Michaelis-Menten datasets are obtained from single time-courses: with gradients over 60 distinct substrate concentrations up to ten times more datapoints per Michaelis-Menten plot than with other systems (see Tables S1-3, Supporting Information) are collected, thus allowing for reliable non-linear fits, so that different kinetic models can be probed, e.g. requiring additional fitting terms for product/substrate inhibition or cooperativity between different substrates. Here, the parallel detection of droplet absorbance in up to twelve points is enabled by the implementation of a rapid and sensitive line scan camera. A new analysis strategy (see Section S1, Supporting Information) identifies droplet boundaries by using the moving average of the signal to prevent incorrect droplet identification. The gradients detected are then fitted to the equation for classic second order rate reactions. The consequence of these improvements is an increase in throughput by an order of magnitude, improved from one to twelve kinetic datasets obtained in 30 minutes (corresponding to 8640 data points per hour) as well as more accurate fits, higher robustness, and shorter time to analysis from raw datasets to Michaelis-Menten plots.

The demand for rapid quantitative analysis systems for the determination of enzyme kinetics with minimal sample consumption (i.e. small reaction volumes and minimal unused dead volumes) is growing,^35^ as the number of quantitatively characterized enzymes is increasingly dwarfed by the emerging sequence data.^3, 5^ The automation and massive scale-down in microfluidic methods provide a solution to this problem.^8, 29-30^ However, several existing microfluidic systems are limited in their throughput as they operate kinetic measurements in a “one enzyme at a time” manner.^24, 26-27, 29, 31, 36-38^ While the data quality in systems that measure concentrations in a one-by-one process is high due to the averaging of measurements from many droplets with the same concentration, these systems have to be reset, cleaned and equilibrated for each new enzyme or variant, compromising throughput.^24, 27, 29, 37^ Systems relying on continuous flows also often have a large overall consumption of precious reagents. Substrate gradients can be generated by variation of substrate and diluent flow rates,^39^ from laminar co-flow on chip,^25^ in capillaries,^40-42^ or by merging droplets in an array system.^38^ Setting up gradients with one droplet representing one concentration is more resource efficient, but the practical solutions can be technically complex. For example, many systems either require expensive robotics^41, 43^ or sophisticated multilayer microfluidic chips with valves that require expertise in fabrication and operation.^26, 36, 38, 44^ The system of Markin et al. (2021), the most comprehensive analysis tool to date, achieves high precision and high throughput (of up to 1,500 enzyme variants in parallel) but the reliance on valves complicates operation and the dependence on a fusion protein, *in vitro* expression and a fluorogenic assay may limit the convenience of its use,^8^ leaving room for versatile systems such as the one described here, despite their lower throughput.

Our platform is comparatively simple: requiring only tubing, pipette tips and laser-cut adapters for the microfluidic part, no chips or specialized, expensive equipment are needed beyond the line scan camera (<1000 USD). Despite this practical simplicity, the number of data points generated is large, while previous systems generated only few data points per kinetic set, which can lead to less accurate non-linear fits.^26, 44, 46-47^ The optical detection unit is easy to assemble, adaptable to any wavelengths in the UV-Vis spectrum and can be flexibly adjusted for custom time intervals in between successive measurements. Examples in the literature often rely exclusively on a fluorescence readout,^8, 24, 29, 36, 38, 41^ which limits the available assays, require custom-made fluorogenic substrates (with non-natural and activated leaving groups) and fluorophores can impose problems by interaction with PDMS.^8, 29^ By contrast our parallel droplet-on-demand platform has been designed around the widely used absorbance readout. Thus, substrates with a wide range of colored leaving groups can be kinetically characterized by exchanging a single LED. This makes the system easy to adjust to a new target reaction, whereas other absorbance-based microfluidic setups only allow for one reaction to be measured at a time and require the exchange and calibration of several LEDs.^27^ Taken together, the combination of miniaturized reaction vessels with multiplexed, high-speed spectrophotometric analyses paves the way for large-scale, semi-automated functional characterization of enzyme mutants and panels of candidate substrates or enzymes.

## Acknowledgments

This research was funded by the Biotechnology and Biological Sciences Research Council (BB/T003545/1), the EPSRC (EP/H046593/1) and the EU H2020 project MetaFluidics (685474). SN received a PhD studentship from AstraZeneca. FH is an ERC Advanced Investigator (695669). The authors thank Timo Kohler for taking photographs of the microfluidic setup; Tomas Buryska, Zbynek Prokop and Jiri Damborsky for useful discussions.

