## Supplementary Information for "High throughput steady-state enzyme kinetics measured in a parallel droplet generation and absorbance detection platform"

**for**

**Table of Contents**

S1. Data analysis to extract Michaelis-Menten kinetics from line camera data 2
S2. Determination of the linear range of the absorbance readout in droplets 4
S3. Supporting Figures 5
S4. Supporting Tables 14
S5. Supporting References 17

### S1. Data analysis to extract Michaelis-Menten kinetics from line camera data

The raw data read from the line camera as droplets pass through each detection point was converted into Michaelis-Menten kinetics plots using the following steps (https://github.com/fhlab/Line_detector_kinetics):

I. The time corresponding to end points of every gradient (i.e. time tag for the final droplet in the gradient), number of droplets composing the gradient and overall gradient duration per detection point were entered manually. Although this step could be automated, it was found to be helpful in the following cases:

- Imperfect monodispersity at the end of gradients (after transfer of the droplet maker to an oil well) leading to some larger droplets to not be counted.
- Two droplets getting too close within the tubing and identified as a single drop, altering the overall droplet number.
- Small differences seen between actual gradient duration (30 seconds) and duration read during the measurements due to local flow rate fluctuations, typically ±1 second.

II. Based on the input gradient times, droplet boundaries were identified from variations in the standard deviation of the moving average of the signal. By using the moving average, sudden signal spikes at the water-oil interfaces (i.e. spikes going above and below the signal corresponding to the actual droplet) are filtered out. This prevents incorrect droplet identification when the signal of the gradient is not monotonic due to the water-oil transitions.

III. After subtraction of the enzyme-only signal baseline, only points belonging to the gradient droplets were analyzed. Every gradient was fit using the following equation, as expected for second-order kinetics reactions:

$$y = ae^{b(t+c)} + d \quad (1)$$

Where $t$ is time, $y$ is the signal, and $a$, $b$, $c$ and $d$ are fitting constants.

IV. The software segmented the fit into the number of droplets counted in step I. This can be done as droplets are generated at fixed rate during substrate infusion. This step circumvented issues arising when droplets accidentally got too close, leading to mismatches in tracking every individual reactor. Time tags were assigned by evaluating the mean time of every segment.

V. All the points corresponding to the same $n^{th}$ droplets were connected and slopes extracted (i.e. reaction rates; reversing the droplet order every other gradient due to the reversal of flow direction).

VI. Enzyme concentration and substrate concentrations were inferred from the infusion/extraction flow rates and initial concentrations as described in a previous study[1] and the Michaelis-Menten parameters $k_{cat}$ and $K_M$ were obtained.

### S2. Determination of the linear range of the absorbance readout in droplets

We performed a linear data regression to determine the linear region of the calibration gradients, considering that the error in concentration can be assumed as a maximum time shift of 1 s between the estimated and actual start of the gradients (Figure S1). The resulting calibration curve indicated that linearity was observed up to just above 1 mM *p*NP.

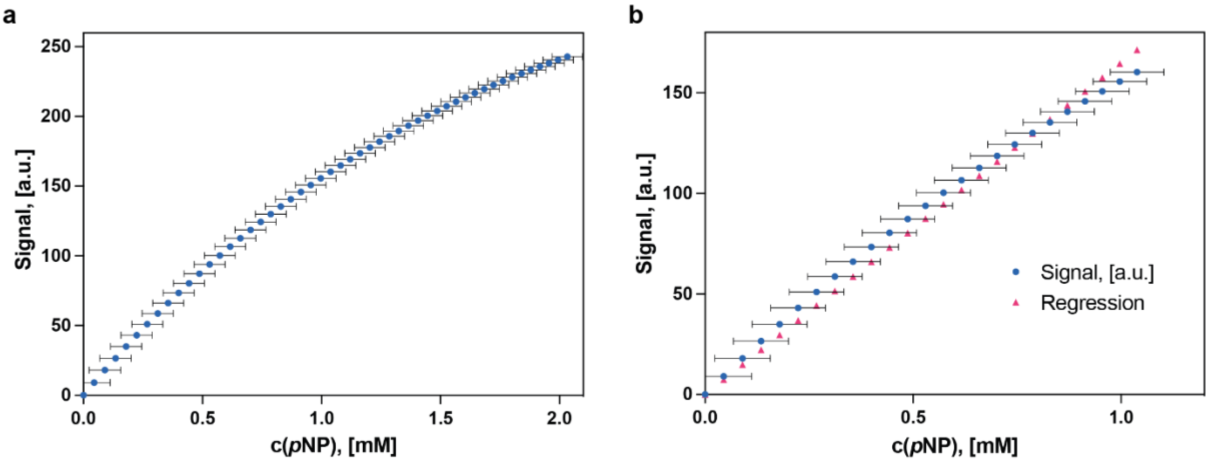


**Figure S1: Calibration to determine the linear range of the detector**. (**a)** Using a model *p*NP gradient, we obtained readings ranging from 0 to 2 mM *p*NP. **(b)** A linear regression confirms good linearity ($R^2$ > 0.99) up to 1 mM *p*NP. The error bars indicate the error in concentration resulting from a 1 s error range in time assignment. The example is shown for detection point 1 but is consistent across all 12 detection points.

After extracting the slope for the linear region, we obtained the limit of detection (LOD) for each detection point by calculating the standard deviation of the filtered signal level for the enzyme-only baseline. Using a confidence level of 95% (2 sigma above the determined background signal), we deduced that the LOD for every detection point ranged from ~5 to 18 µM (Figure S2).

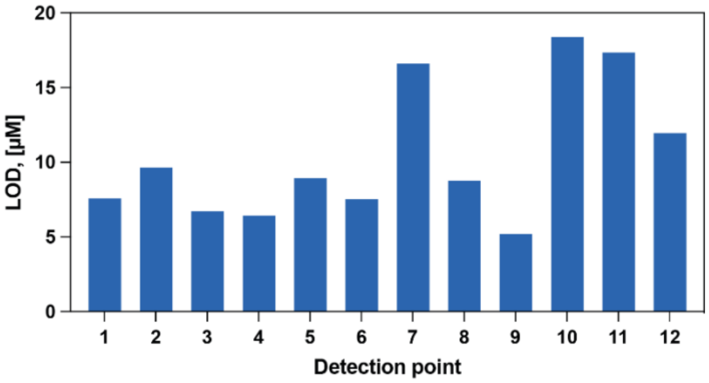


**Figure S2**. **Typical limits of detection for *p*-nitrophenol at the 12 detection points.** We ascribe the variations in LOD to imperfect incoming light power uniformity, and the imprecise alignment of tubing with respect to the threading holes, through which light propagates.

### S3. Supporting Figures

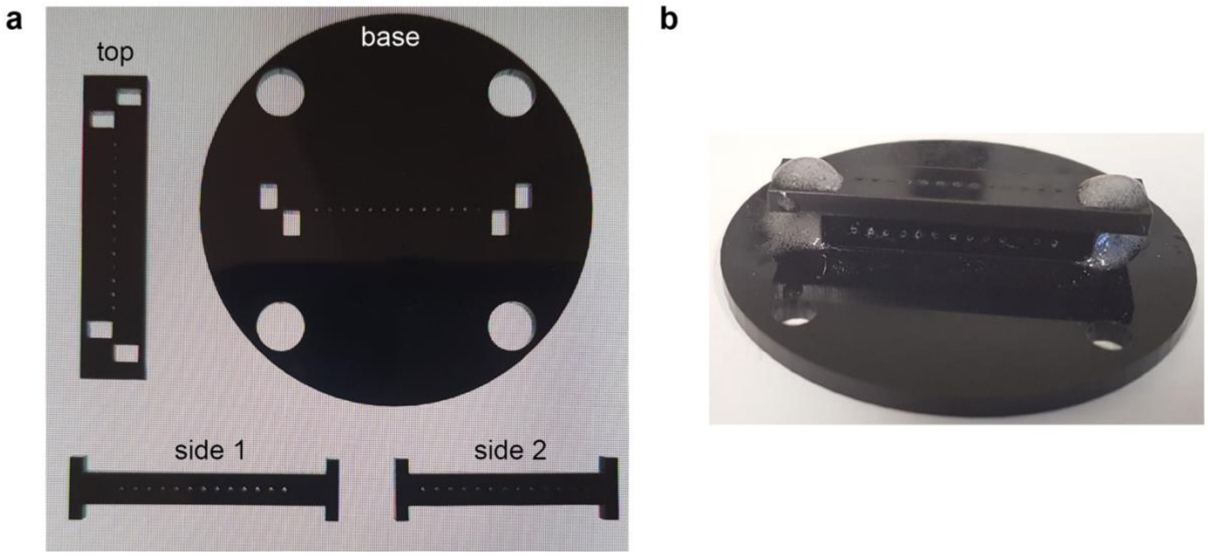


**Figure S3. Fluidic connector aligning tubing for detection by the line camera. (a)** Individual components made of black acrylic. The base, top and two side building blocks have evenly spaced holes (13 of which only 12 are used) that align with each other when assembled. **(b)** Assembled fluidic connector. The adaptor is placed on the line camera (see Figure 2, main article) and light from the collimated light sheet passes vertically through the holes from the top. Tubing is inserted horizontally through the holes in the two vertical side walls and pass through the light path.

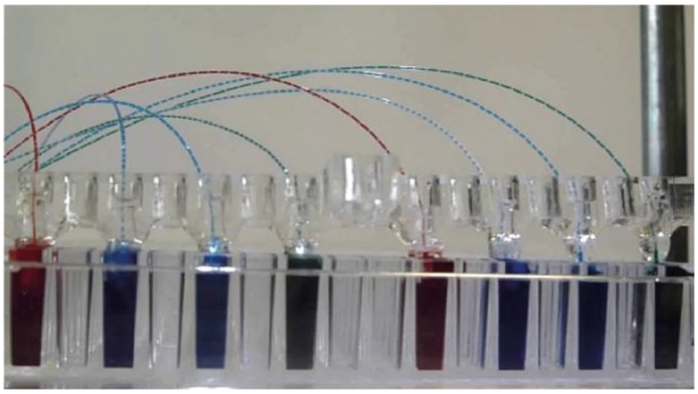

**Figure S4. Towards automated sampling from microtiter plates**. In this example, eight droplet generators operate in parallel, such that eight enzymatic reactions could be probed at the same time. The fluidic adaptor was made up of a main slab in clear acrylic with eight holes ~ 5 mm diameter, aligned with every other well of a 384-well plate. In each hole, the cut end of a gel loading tip was glued in its center. After insertion of the ultra-microbore PTFE tubing in the loading tip, oil can be pipetted into the sealed holes. A multi-rack syringe pump operates in withdrawal mode at constant flow rate across the eight fluidic lines.

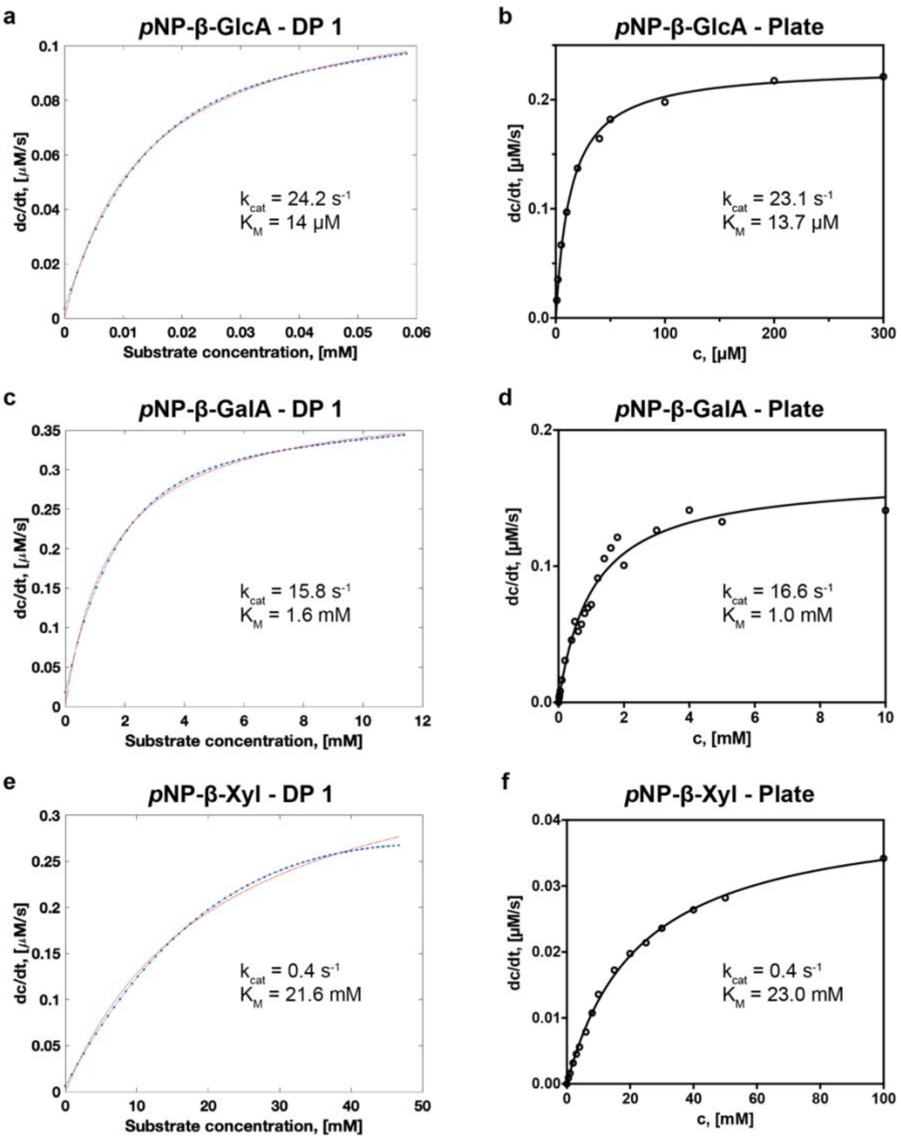


**Figure S5: Side-by-side comparison of Michaelis-Menten kinetics determined from in-droplet measurements (for selected detection points) and microtiter plate measurements – part 1.** Kinetic data were obtained for SN243 with *p*NP-β-GlcA **(a,b)**, *p*NP-β-GalA **(c,d)** and *p*NP-β-Xyl **(e,f)**. Resulting plots from absorbance detection at 405 nm in droplets **(a,c,e)** and in microtiter plates **(b,d,f)** were compared. DP: detection point.
Note: The differences in initial velocities (given as dc/dt in [µM/s]) between the two assay formats result from the usage of different enzyme concentrations. The microtiter plate data were obtained from Neun et al. (2022).[2]

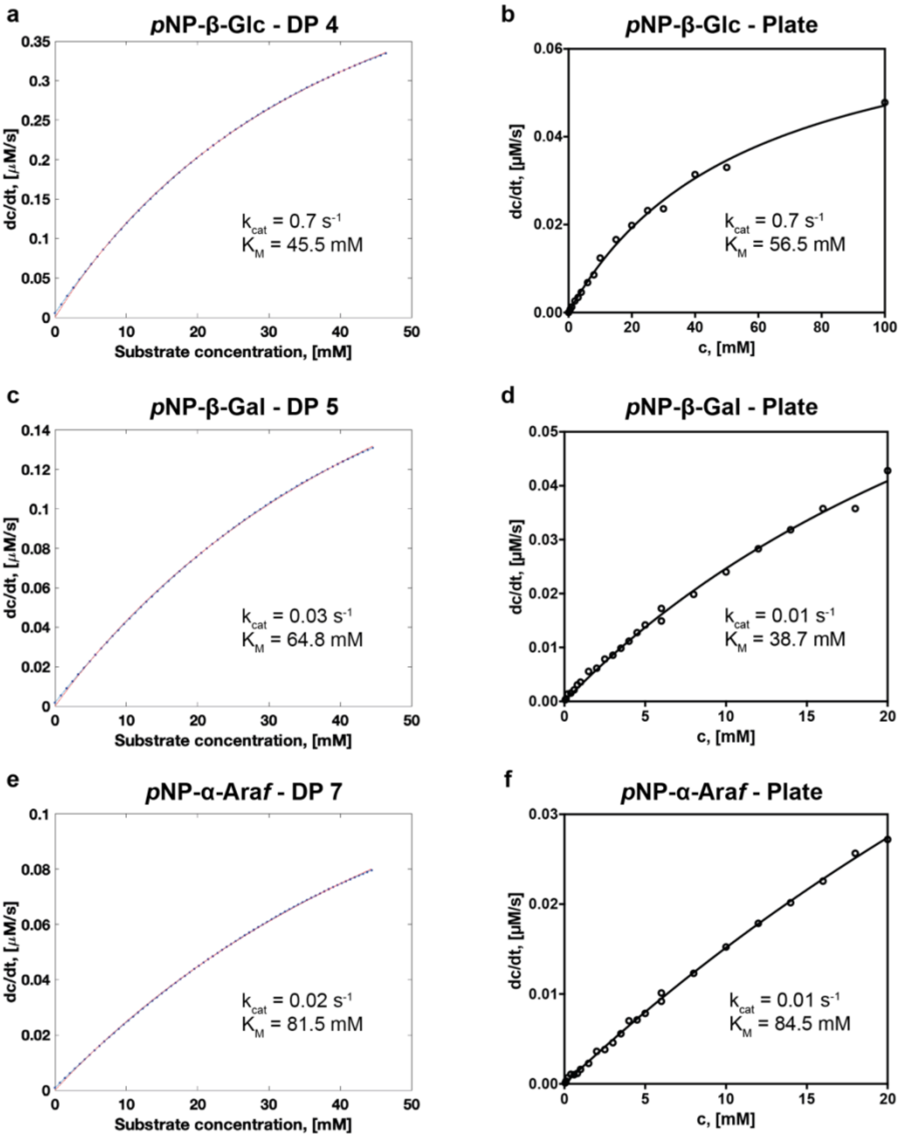


**Figure S6: Side-by-side comparison of Michaelis-Menten kinetics determined from droplet (for selected detection points) and microtiter plate measurements– part 2.** Kinetic data were obtained for SN243 with *p*NP-β-Glc **(a,b)**, *p*NP-β-Gal **(c,d)** and *p*NP-α-Ara*f* **(e,f)**. Resulting plots from absorbance detection at 405 nm in droplets **(a,c,e)** and in microtiter plates **(b,d,f)** were compared. DP: detection point.

Note: The differences in initial velocities (given as dc/dt in [µM/s]) between the two assay formats result from the usage of different enzyme concentrations. The microtiter plate data were obtained from Neun et al. (2022).[2]

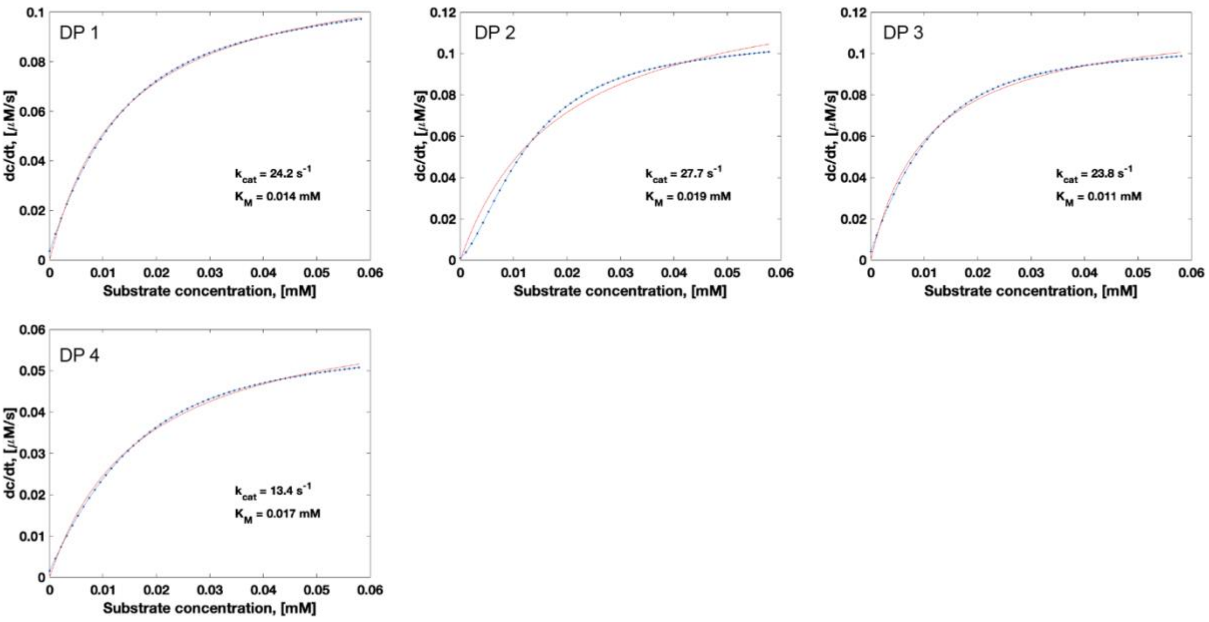


**Figure S7: Individual plots for 4 detection points measured for the same droplet gradient for the determination of Michaelis-Menten kinetics of SN243 with *p*NP-β-GlcA.** Initial velocities for the reaction extrapolated from droplet gradient measurements (blue) and fitting functions (red) are plotted for all detection points used to determine the average parameters indicated in Table 1 of the main manuscript. DP: detection point.

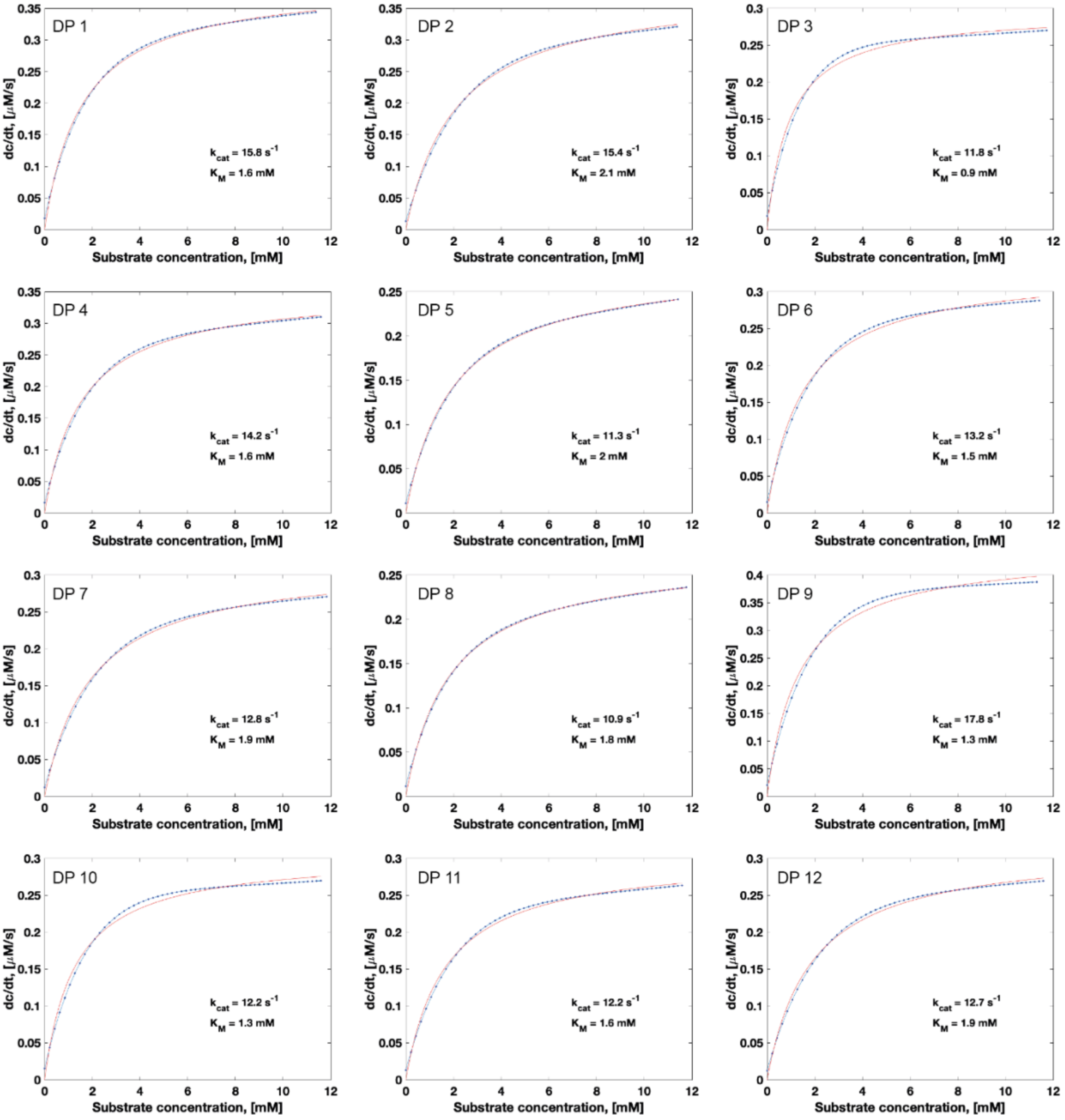


**Figure S8: Individual plots for the 12 detection points for the accurate determination of Michaelis-Menten kinetics of SN243 with *p*NP-β-GalA in a single experiment.** Detection points 1-4, 5-8 and 9-12 correspond to three distinct substrate concentration gradients. Initial velocities for the reaction extrapolated from droplet gradient measurements (blue) and fitting functions (red) are plotted for all detection points used to determine the average parameters indicated in Table 1 of the main manuscript. DP: detection point.

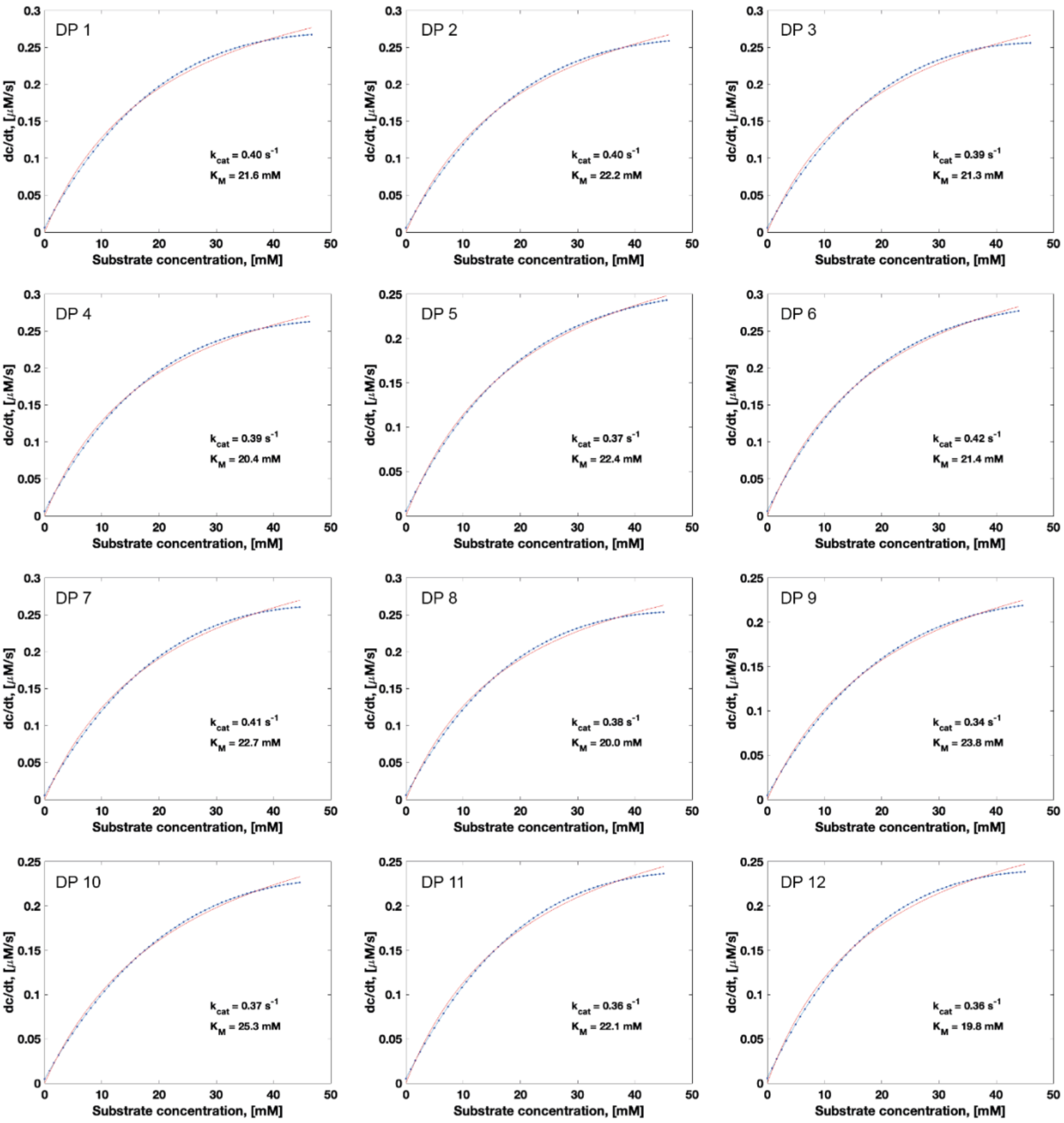


**Figure S9: Individual plots for 12 detection points for the accurate determination of Michaelis-Menten kinetics of SN243 with *p*NP-β-Xyl in a single experiment.** Detection points 1-4, 5-8 and 9-12 correspond to three distinct substrate concentration gradients. Initial velocities for the reaction extrapolated from droplet gradient measurements (blue) and fitting functions (red) are plotted for all detection points used to determine the average parameters indicated in Table 1 of the main manuscript. DP: detection point.

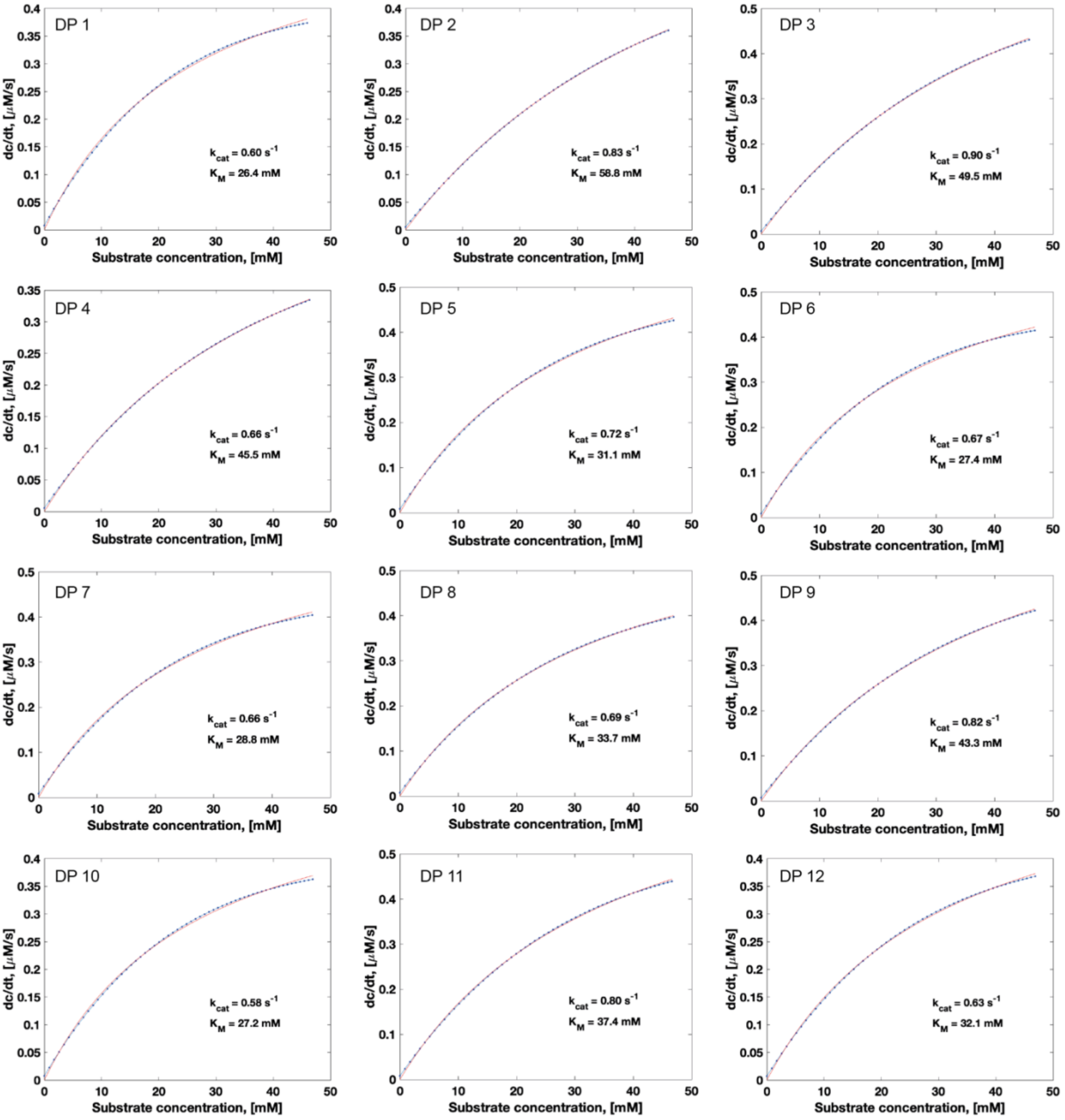


**Figure S10: Individual plots for 12 detection points for the determination of Michaelis-Menten kinetics of SN243 with *p*NP-β-Glc in a single experiment.** Detection points 1-4, 5-8 and 9-12 correspond to three distinct substrate concentration gradients. Initial velocities for the reaction extrapolated from droplet gradient measurements (blue) and fitting functions (red) are plotted for all detection points used to determine the average parameters indicated in Table 1 of the main manuscript. DP: detection point.

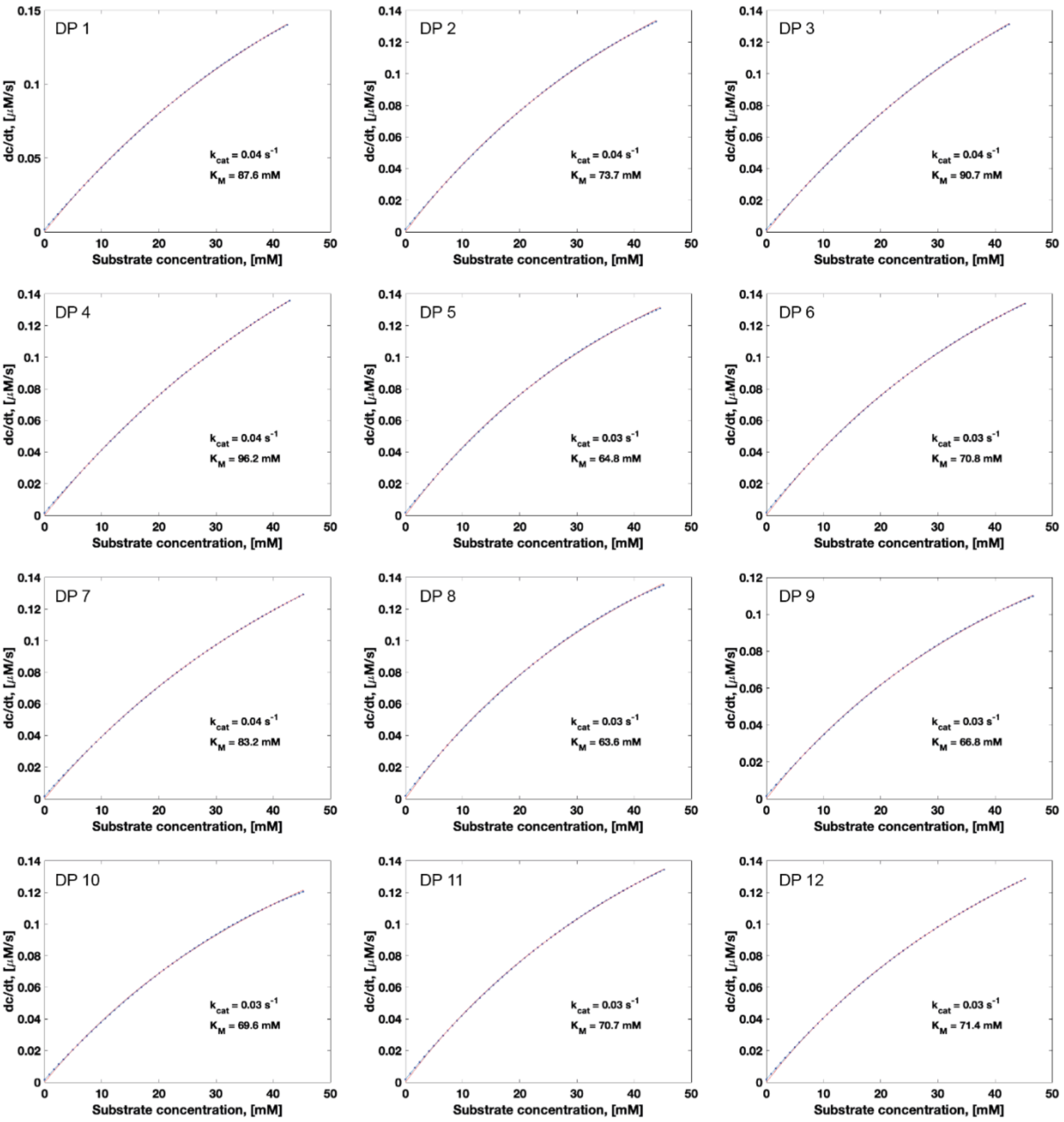


**Figure S11: Individual plots for 12 detection points for the determination of Michaelis-Menten kinetics of SN243 with *p*NP-β-Gal in a single experiment.** Detection points 1-4, 5-8 and 9-12 correspond to three distinct substrate concentration gradients. Initial velocities for the reaction extrapolated from droplet gradient measurements (blue) and fitting functions (red) are plotted for all detection points used to determine the average parameters indicated in Table 1 of the main manuscript. DP: detection point.

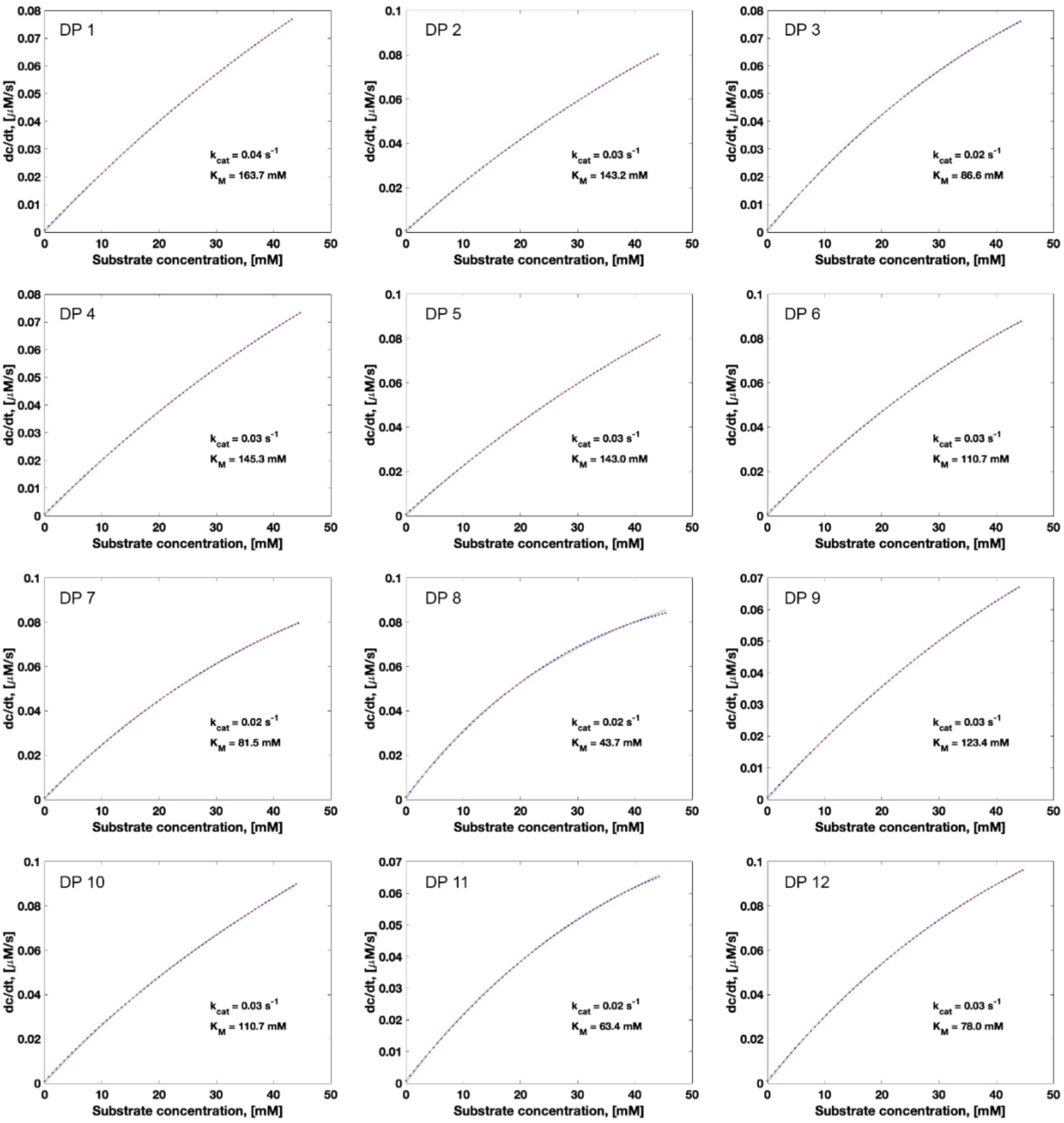


**Figure S12: Individual plots for 12 detection points for the determination of Michaelis-Menten kinetics of SN243 with *p*NP-α-Ara*f* in a single experiment.** Detection points 1-4, 5-8 and 9-12 correspond to three distinct substrate concentration gradients. Initial velocities for the reaction extrapolated from droplet gradient measurements (blue) and fitting functions (red) are plotted for all detection points used to determine the average parameters indicated in Table 1 of the main manuscript. DP: detection point.

### S4. Supporting Tables

**Table S1: Microfluidic systems for kinetic analysis using droplets generated and measured in continuous flow devices.**

| Ref. | Concentrations one-by-one or gradient? | Readout | Different reactions in parallel | Individual droplets or averaging over many droplets? | Time points from same or separate experiment | Datapoints per kinetic dataset | Datapoints per $v_0$ determination | Duration of operation | Reaction time scale | Kinetic analysis | Difficulty |
|---|---|---|---|---|---|---|---|---|---|---|---|
| [3] | One-by-one | Fluorescence intensity | 1 | Averaging | Same | 4 | ~10 | min-h | Fast | Pre-steady state analysis | ++ |
| [4] | | | | | | 10-15 | | | | | |
| [5-6] | | Electrochemistry | | | Separate | 5 | | | | | +++ |
| [7] | Gradient (4 concentrations by laminar flow) | Fluorescence intensity | | | Same | 10 | | | Slow | Steady-state analysis | +++ |
| [8] | One-by-one | Absorbance (LEDs) | | | | 6 | 7 | | | | ++ |
| [9] | | Photothermal interferometry | | | | 5 | 4 | | | | |
| [10] | gradient (by flow rate variation) | Fluorescence intensity | | Individually | | ~$10^3$ | 35 | | Fast | | |
| [11] | Inhibitor gradient (generated in capillary prior to droplet formation) | Electrochemistry | | | n.a. | 40-50 | 1 | | Slow | Inhibitor potency measurement | +++ |
| [12] | One-by-one | Absorbance optical fiber-based) | | Averaging | | 7 | | | | | ++ |
| [13] | Inhibitor gradient (generated in capillary prior to droplet formation) | Laser-induced fluorescence | | | | 28 | | | | | ++ |

*(low concentrations/linear range of Michaelis-Menten plot not captured); n.a.: not applicable.

**Table S2: Microfluidic systems for kinetic analysis using droplets in segmented flow.**

<table>
<tr><th>Ref.</th><th>Concentrations one-by-one or gradient?</th><th>Readout</th><th>Different reactions in parallel</th><th>Individual droplets or averaging over many droplets?</th><th>Time points from same or separate experiment</th><th>Datapoints per kinetic dataset</th><th>Datapoints per $v_0$ determination</th><th>Duration of operation</th><th>Reaction time scale</th><th>Kinetic analysis</th><th>Difficulty</th></tr>
<tr><td>[14]</td><td>Inhibitor gradient in droplets-on-demand (pre-pipetted microtiter plate)</td><td rowspan="2">Laser-induced fluorescence</td><td rowspan="3">1</td><td>Individually</td><td rowspan="9">Same</td><td>7</td><td>7</td><td>hours</td><td rowspan="5">Slow</td><td rowspan="3">Inhibitor potency measurement</td><td rowspan="3">+++</td></tr>
<tr><td>[15]</td><td>Inhibitor gradient (5 concentrations by flow ratio adjustment)</td><td>Averaging</td><td>5 substrate and 5 inhibitor concentrations</td><td>6</td><td rowspan="2">min-h</td></tr>
<tr><td>[16]</td><td>gradient in droplets-on-demand (by coalescing defined numbers of reagent droplets)</td><td rowspan="4">Fluorescence intensity</td><td rowspan="7">Individually</td><td>32</td><td>12</td></tr>
<tr><td>[17]</td><td>concentration gradient in droplets-on-demand (generated in capillary prior to droplet formation)</td><td>102</td><td>24</td><td>6</td><td>hours</td><td>Compound screening</td><td>+++</td></tr>
<tr><td>[18]</td><td>gradient in droplets-on-demand (by valve-based system)</td><td>2</td><td rowspan="2">~5</td><td>10</td><td rowspan="5">min-h</td><td>Steady-state analysis</td><td rowspan="2">+++</td></tr>
<tr><td>[19]</td><td>gradient in droplets-on-demand (6 concentrations from separate syringes)</td><td rowspan="3">1</td><td>20</td><td>Fast & slow</td><td>(Pre-)steady-state analysis</td></tr>
<tr><td>[1]</td><td>concentration gradient in droplets-on-demand (by merging droplets of different concentration and volume ratios)</td><td rowspan="3">Absorbance</td><td>24</td><td>10</td><td rowspan="3">Slow</td><td rowspan="3">Steady-state analysis</td><td rowspan="3">+</td></tr>
<tr><td>[20]</td><td rowspan="2">Concentration gradient in droplets-on-demand (droplets generated while changing concentration in source well)</td><td>150</td><td rowspan="2">6</td></tr>
<tr><td>This study</td><td>12</td><td>60</td></tr>
</table>

**Table S3: Droplet-free microfluidic systems for kinetic analysis.**

<table>
<tr><th>Ref.</th><th>Concentrations one-by-one or gradient?</th><th>Readout</th><th>Different reactions in parallel</th><th>Individual measurements or averaging over time?</th><th>Time points from same or separate experiment</th><th>Datapoints per kinetic dataset</th><th>Datapoints per $v_0$ determination</th><th>Duration of operation</th><th>Reaction time scale</th><th>Kinetic analysis</th><th>Difficulty</th></tr>
<tr><td>[21]</td><td rowspan="3">One-by-one</td><td>Laser-induced fluorescence</td><td rowspan="6">1</td><td rowspan="3">Averaging</td><td>Same</td><td rowspan="3"><10</td><td>15-20</td><td rowspan="2">min-h</td><td rowspan="7">slow</td><td>Steady-state analysis</td><td>++</td></tr>
<tr><td>[22]</td><td>Fluorescence intensity</td><td rowspan="2">Separate</td><td rowspan="2">~5</td><td rowspan="2">Reuse of enzyme for steady-state analysis</td><td rowspan="2">++</td></tr>
<tr><td>[23]</td><td>Electrochemistry</td><td rowspan="2">hours</td></tr>
<tr><td>[24]</td><td>Concentration gradient (in microprocessors)</td><td rowspan="2">Fluorescence intensity</td><td rowspan="2">Individually</td><td rowspan="4">Same</td><td>11</td><td>~30</td><td rowspan="3">Steady-state analysis</td><td>++++</td></tr>
<tr><td>[25]</td><td rowspan="3">One-by-one</td><td>12</td><td>60</td><td>min-h</td><td>+</td></tr>
<tr><td>[26]</td><td>Infrared</td><td>Averaging</td><td>9</td><td>8</td><td>hours</td><td>+++</td></tr>
<tr><td>[27]</td><td>Fluorescence intensity</td><td>1500</td><td>Individually</td><td>10-15</td><td>15-20</td><td>days</td><td>Expression, purification and steady-state analysis on one chip</td><td>++++</td></tr>
</table>
